# Rapid modeling of an ultra-rare epilepsy patient variant in mice by in utero prime editing

**DOI:** 10.1101/2023.12.06.570164

**Authors:** Colin D. Robertson, Patrick Davis, Ryan R. Richardson, Philip H. Iffland, Daiana C. O. Vieira, Marilyn Steyert, Paige N. McKeon, Andrea J. Romanowski, Garrett Crutcher, Eldin Jašarević, Steffen B. E. Wolff, Brian N. Mathur, Peter B. Crino, Tracy L. Bale, Ivy E. Dick, Alexandros Poulopoulos

**Affiliations:** Department of Pharmacology and UM-MIND, University of Maryland School of Medicine, Baltimore, MD, USA; Department of Neurology, Boston Children’s Hospital, Harvard Medical School, Boston, MA, USA; Department of Neurology, and UM-MIND, University of Maryland School of Medicine, Baltimore, MD, USA; Department of Physiology and UM-MIND, University of Maryland School of Medicine, Baltimore, MD, USA

## Abstract

Generating animal models that mirror a patient’s seizures within clinically-useful timeframes is an important step toward advancing precision medicine for genetic epilepsies. Here we report a somatic cell genome editing approach that rapidly incorporated a patient’s genomic variant into mice, which developed seizures recapitulating elements of the patient’s pathology. This approach offers a versatile in vivo platform for clinical, preclinical, and basic research applications, including tailoring pharmacotherapy, assessing variants of uncertain significance, and screening compounds to develop drugs for rare epilepsies. As proof-of-principle, we modeled an epilepsy patient with an ultra-rare variant of the NMDA receptor subunit GRIN2A using prime editing in utero directly in the developing brain of wild-type mice. This methodology achieved high-fidelity genome editing in vivo sufficient to induce frequent spontaneous seizures without necessitating germline modification or extensive breeding. Leveraging the speed and versatility of this approach, we propose a generalizable workflow to generate bedside-to-bench animal models of individual patients within weeks. This advance holds promise for providing a cost-effective, expedient in vivo testing platform that reduces barriers to access for precision medicine, and accelerates drug development for rare and neglected neurological conditions.

---

Genetic epilepsies afflict nearly 1 in 2000 children born^1^. They are exceptionally heterogeneous in etiology, clinical presentation, and responsiveness to treatment^2,3^. With nearly 1000 associated gene variants already identified, they comprise a collection of ultra-rare diseases^4^ for which tailored treatment options are limited due to challenges in assembling large study trial cohorts.

Without a systematic path for tailoring pharmacotherapy, over half of all epilepsy patients are faced with the burden of testing trial medications on themselves, and 30% nonetheless remain unresponsive to their current treatment^5^. Even in patients with known genetic causes of epilepsy, reliable prediction of therapeutic or deleterious response to medication trials remains elusive^3^. In this space of inadequately treated ultra-rare epilepsies, a platform to identify patient-specific efficacies by screening existing anti-epileptics or off-label use of compounds approved for humans would offer a path toward systematizing treatment selection in a manner otherwise not clinically feasible.

Precision medicine approaches hold great promise for enabling tailored therapies for rare-variant conditions. Genetic engineering and genome editing technologies enable the introduction of patient gene variants into transgenic mice, to serve as animal models in which treatments are assessed. However, due to the laborious, costly, and time-consuming process to generate mouse lines through breeding for germline transmission, only a fraction of patient variants has been developed into animal models. This bottleneck in the experimental pipeline has resulted in the available animal strains becoming singular in vivo models of the disease as a whole, rather than models of the specific variants, further limiting our understanding and treatment of these diverse conditions^3,6^. This approach is further restricted in producing inbred, genetically homogenous animal lines that do not recapitulate the phenotypic variability seen in human pedigrees of genetic epilepsy^7,8^.

In order for personalized animal models to become a precision medicine tool that can be applied broadly in a clinical setting, the technology to generate animals harboring individual patient variants must be i) rapid; applicable in clinically-relevant timescales, ii) versatile; applicable to a variety of gene variants, and iii) validatable; able to recapitulate identifiable features of the individual’s clinical presentation to be measured against therapeutic interventions.

We present here an experimental approach which demonstrates these features by leveraging developments in somatic cell genome editing and new precision editors^9–11^ to introduce patient variants into the genome of relevant cell types without the need for breeding transgenic animal strains. By performing somatic cell genome editing directly in the brain in vivo, our workflow circumvents germline transmission and the requirement for breeding, resulting in ready-to-use animal models within weeks (Fig. 1).

**Fig. 1.**
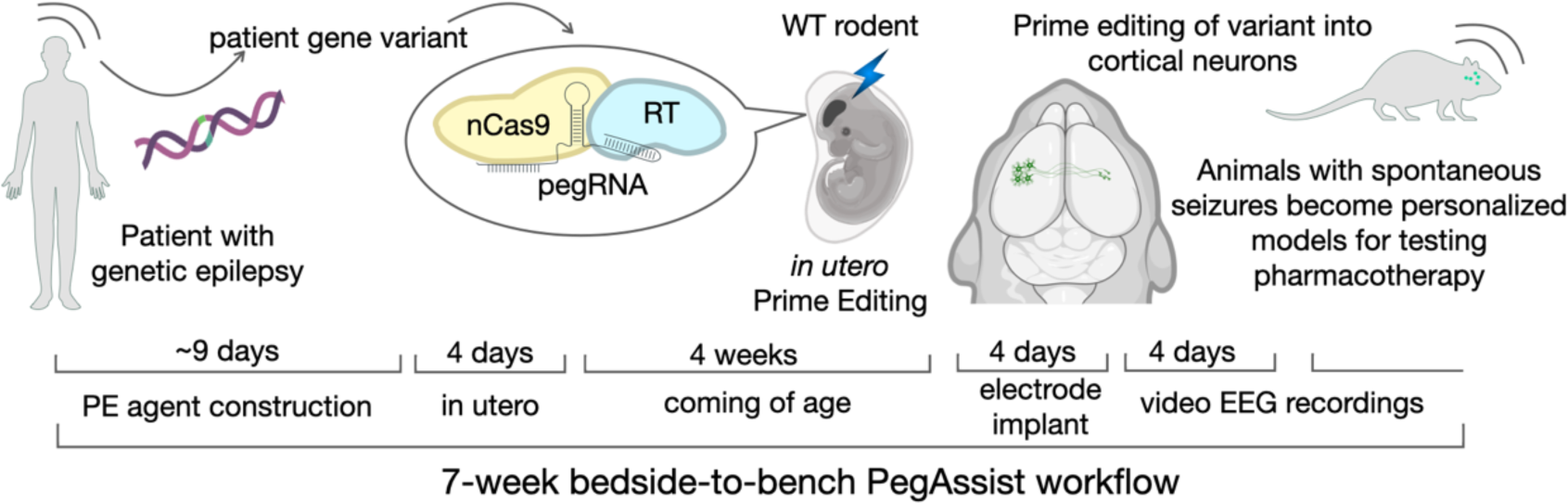
In utero prime editing workflow for generating personalized animal models. Schema depicting the 7-week workflow for personalized animal models, from variant identification, to editing agent construction, in utero delivery, and baseline analysis to identify animal for personalized models.

We demonstrate proof-of-principle somatic cell genome editing of an epilepsy patient point-variant directly into neurons of the cerebral cortex of wild-type mice. The resulting mice displayed frequent, spontaneous seizures reproducing several core characteristics of the clinical presentation of the patient. To our knowledge, this is the first demonstration of prime editing performed in vivo specifically targeting neurons, and the first proof-of-principle demonstration of a patient-specific, neurogenetic disease model in wild-type mice. This platform has the potential to be generalizable across a range of genetic variants, and increases the time- and cost-effectiveness of animal modeling to enable use in clinical settings for tailoring treatment options to individual patients.

## Prime editing 3b demonstrates high editing precision

In vivo somatic cell genome editing requires an editing agent with high on-target precision, i.e. a high ratio of correct versus incorrect edits on the target locus^12^. This metric is the major determinant of an editor’s signal-to-noise ratio, and the key limiting factor for in vivo use, given that without ex vivo selection, both intended and unintended edits will persist and accumulate in the body^10^.

We thus began by screening available and engineered high-performance genome editors^13,14^ for high on-target precision. We used a high-throughput platform to measure and compare on-target precision of different editing agents by point editing a genomically-encoded Blue Fluorescent Protein (BFP) gene to introduce a H62Y substitution, which corresponds to the sequence for Green Fluorescent Protein (GFP). Precise editing would convert BFP to GFP, while imprecise loss-of-function edits (e.g.: indels) would result in loss of fluorescence (Fig. 2a).

**Fig. 2.**
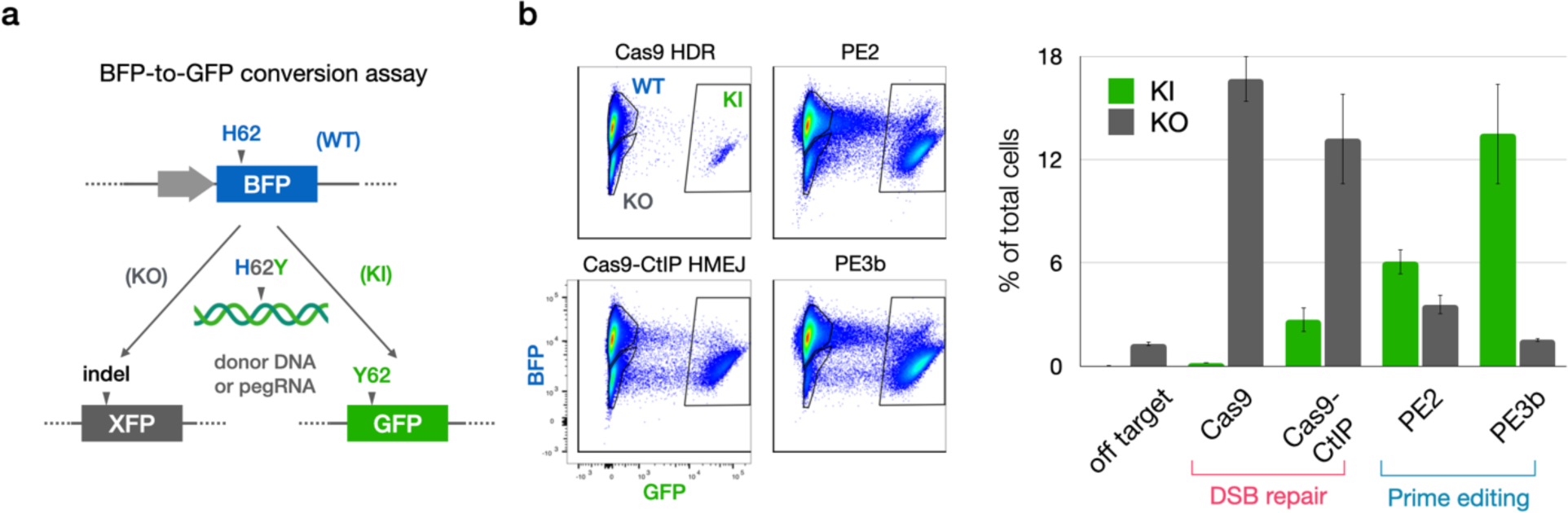
Screening editing agents for precision identifies PE3b as most precise editor. **a**, Schema of BFP-to-GFP conversion assay to assess editor precision. Cells with genomically encoded BFP were transfected with editing agents to introduce H62Y edit, which corresponds to the sequence of GFP. Precise editing converts cells from blue to green, while imprecise edits cause loss of fluorescence. Editing precision is calculated as the ratio of green/dark cells. **b**, FACS plots and quantification of blue unedited (WT), green correctly edited (knock-in; KI), and dark incorrectly edited (knock-out; KO) HEK cells after treatment with double-strand-break (DSB) repair editors Cas9 (with homology directed recombination [HDR] template) and Cas9-CtIP (with homology-mediated end-joining [HMEJ] template), and prime editing strategies PE2 and PE3b. PE3b outperforms other editing strategies through both higher KI rates and lower KO rates, providing the highest efficiency and precision.

With this strategy we quantified editing precision of homology-directed recombination (HDR) and homology-mediated end-joining (HMEJ) strategies with Cas9 and Cas9-CtIP^13^, as well as of reverse transcriptase-mediated editing with Prime Editor (PE) and PE fused to hRad51 in both PE2 and PE3b strategies^14^. This screen demonstrated exceptionally high on-target precision of point editing with the PE3b strategy, yielding the correct edit over 5-fold more frequently than the aggregate of all other edits (Fig. 2). PE3b has independently been shown to outperform other prime editing strategies ex vivo by minimizing off-target outcomes^15^.

PE is a hybrid ribonucleoprotein consisting of a protein fusion of Cas9 nickase (nCas9) and reverse transcriptase (RT), in complex with a hybrid pegRNA consisting of a “spacer” sequence for nCas9 targeting, a “primer binding sequence” that hybridizes with nicked genomic DNA, and template sequence for RT to encode the edit. To employ the PE3b strategy, an independent gRNA with no RT component directs PE to nick the complementary strand to encourage productive pegRNA-mediated editing (Fig. 3a). To facilitate the design and production of PE agents, we created pegassist.app, a python-based webtool and plasmid set offered through Addgene (Extended Data Fig. 1 and Methods). This webtool may be used in tandem with other pegRNA design tools to optimize editing agent production^16,17^.

**Fig. 3.**
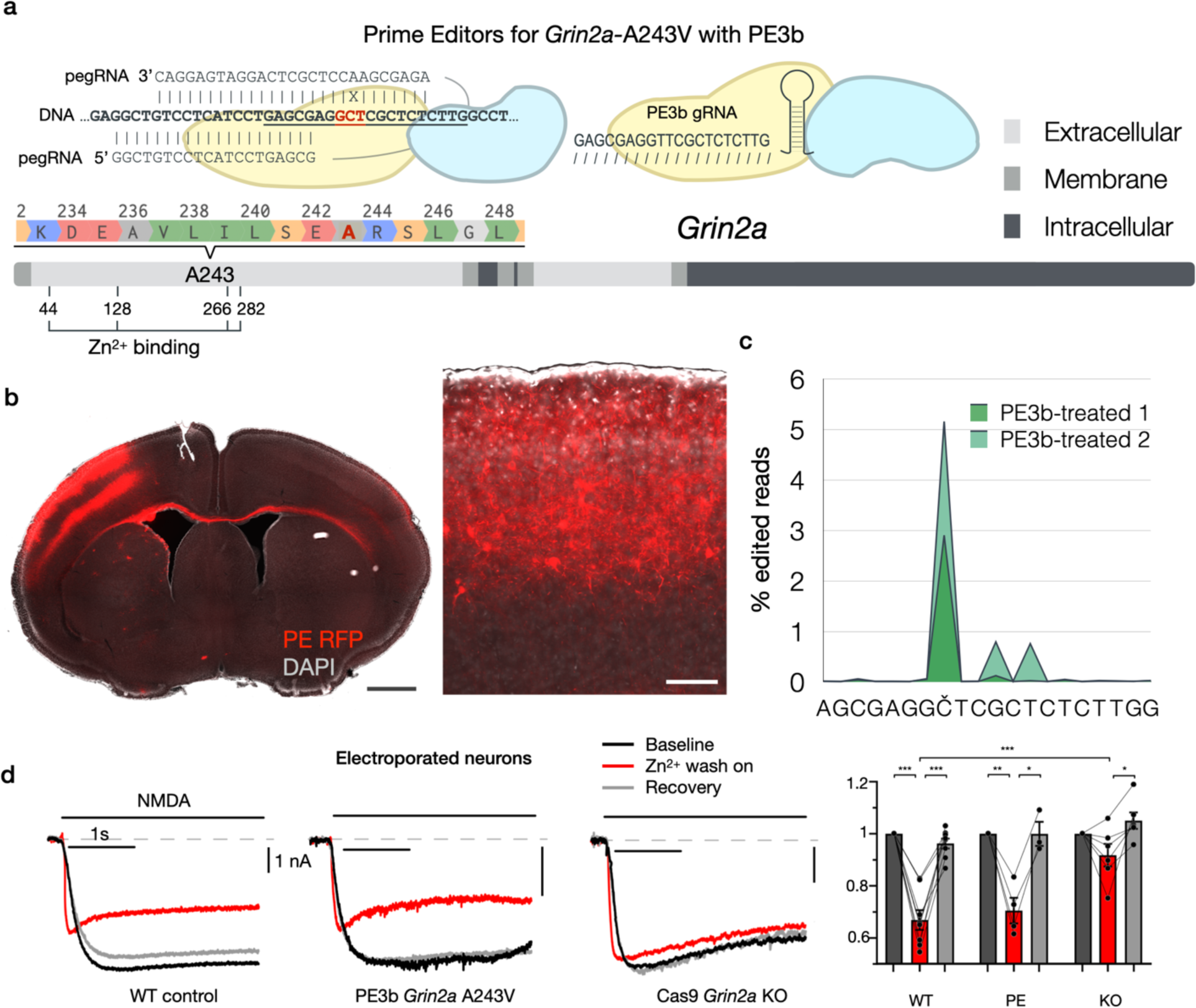
In vivo neuronal genome editing of epilepsy patient variant *GRIN2A*-A243V with in utero prime editing. **a**, Alignment and schema of the WT *Grin2A* target sequence with the pegRNA and PE3b gRNA used to introduce edit A243V. Underlined is the sequence targeted by the PE3b gRNA. In red is the codon for A243. X shows the mismatch between target sequence and pegRNA RT template that introduces the A243V edit. The grey scale bar represents GRIN2A protein primary sequence with cellular topology as indicated in the key. Edited residue A243 and critical residues for Zn^2+^-binding are indicated. **b**, Coronal section of DAPI-stained (grey) brain electroporated with PE and fluorescent marker (red) in centrolateral cortex. Magnified inset shows electroporated upper-layer pyramidal neurons expressing PE. Scale bar = 1 mm, inset 100 μm. **c**, Sequencing of fluorescence-sorted cortical neurons from PE3b-treated mice showing the percentage of sequencing reads that deviated from the reference sequence around the genomic target sequence encoding A243. The intended target nucleotide is marked as Č. Editing predominantly occurs on the intended position. **d**, Example traces and amplitudes of currents recorded from in utero electroporated cultured cortical neurons; baseline (black), Zn^2+^ wash-on (red), recovery (grey); amplitudes normalized to baseline of each sample. Zn^2+^ blockade was detected in WT control (n=8) and PE3b *Grin2a-*A243V cells (n=4), but not Cas9 *Grin2A* KO cells (n=6), indicating that treatment with PE3b does not cause significant loss of function (*p<0.05, ***p<0.001, via students T-test).

## In utero prime editing of GRIN2A variant from epilepsy patient

The exceptional performance of PE3b prompted us to explore its use directly in vivo to model an individual patient variant in wild-type mice. We selected to model a patient with self-limited epilepsy with centrotemporal spikes (SeLECTS) reported with an ultra-rare missense variant, A243V, in the *GRIN2A* gene, encoding the 2A subunit of the N-methyl-D-aspartate (NMDA) type glutamate receptor^18^. GRIN genes are hotspot loci with hundreds of ultra-rare loss- and gain-of-function variants identified to cause conditions, collectively termed GRINopathies, that commonly present with seizures of wide-ranging severity and cognitive comorbidity^19^. Importantly, *Grin2a* knockout mice do not have spontaneous seizures^20^, allowing us to discriminate unintended loss-of-function from intended gain-of-function point-editing at the level of phenotype manifestation.

We constructed PE3b agents using pegassist.app to edit the A243V patient variant into the *Grin2a* locus of the mouse (Fig. 3a). PE and fluorescent reporter plasmids were injected into the lateral telencephalic ventricle of E15 mouse embryos and targeted by in utero electroporation to upper layer pyramidal neurons in centrolateral cortex (Fig. 3b), analogous to the area of centrotemporal cortex, where epileptiform activity is detected in patients with SeLECTS. Animals electroporated in utero with either PE or control plasmids came to term and were allowed to reach adulthood in their home cage.

We directly assessed editing performance in neurons in vivo in two PE3b-treated mice by dissociating and sorting fluorescent cortical neurons from the electroporated target area of cortex. RNA sequenced from sorted cells showed moderate editing efficiency, but high editing precision: the A243V edit was present in ∼5% of reads, while less than 1% of reads displayed any on-target errors (Fig. 3c and Extended Data Fig. 2).

This performance is similar to the >5-fold prevalence of precise edits we observed for PE3b editing in vitro (Fig. 2b), and corresponds to orders-of-magnitude higher precision than other knockin approaches we previously tested^12^. Further, this likely is an underestimate of editing precision, since substitutions appear in sequencing reads as technical artifacts, e.g. due to RT or PCR amplification errors during library preparation^21^, which we did not attempt to discriminate from true editing errors. Importantly, the rate of insertion / deletion events (indels) was minimal (<0.001%), indicating that loss-of-function effects are not a significant editing outcome.

We additionally confirmed using electrophysiology that PE-electroporated neurons do not display *Grin2a* loss of function. NMDA currents of PE-electroporated neurons in culture were largely normal, unlike Cas9-electroporated *Grin2a* knockout neurons, which displayed pronounced reduction in Zn^2+^ blockade of NMDA currents (Fig. 3d), as expected by *Grin2a* loss of function^22^. Using exogenous expression in HEK cells, we corroborated that the *Grin2a*-A243V variant did not measurably alter Zn^2+^ gating (Extended Data Fig. 3), in contrast to a previous report in oocytes^18^. Taken together, these data show that our in utero PE3b strategy successfully incorporated the patient variant into the *Grin2a* locus of neurons in vivo, without detectable loss of function.

## In utero prime edited mice carrying patient variant display spontaneous seizures

Having confirmed in vivo editing in a subset of neurons in centrolateral cortex of wild-type mice, we proceeded to monitor 7 PE-electroporated “PegAssist” (PA) and 6 control-electroporated (CT) mice using video-EEG for 96 hours to determine whether animals present any pathological features associated with SeLECTS. 3 of 7 PA animals displayed spontaneous seizures with electrographic features similar to those seen in SeLECTS patients (Fig. 4)^23,24^.

**Fig. 4.**
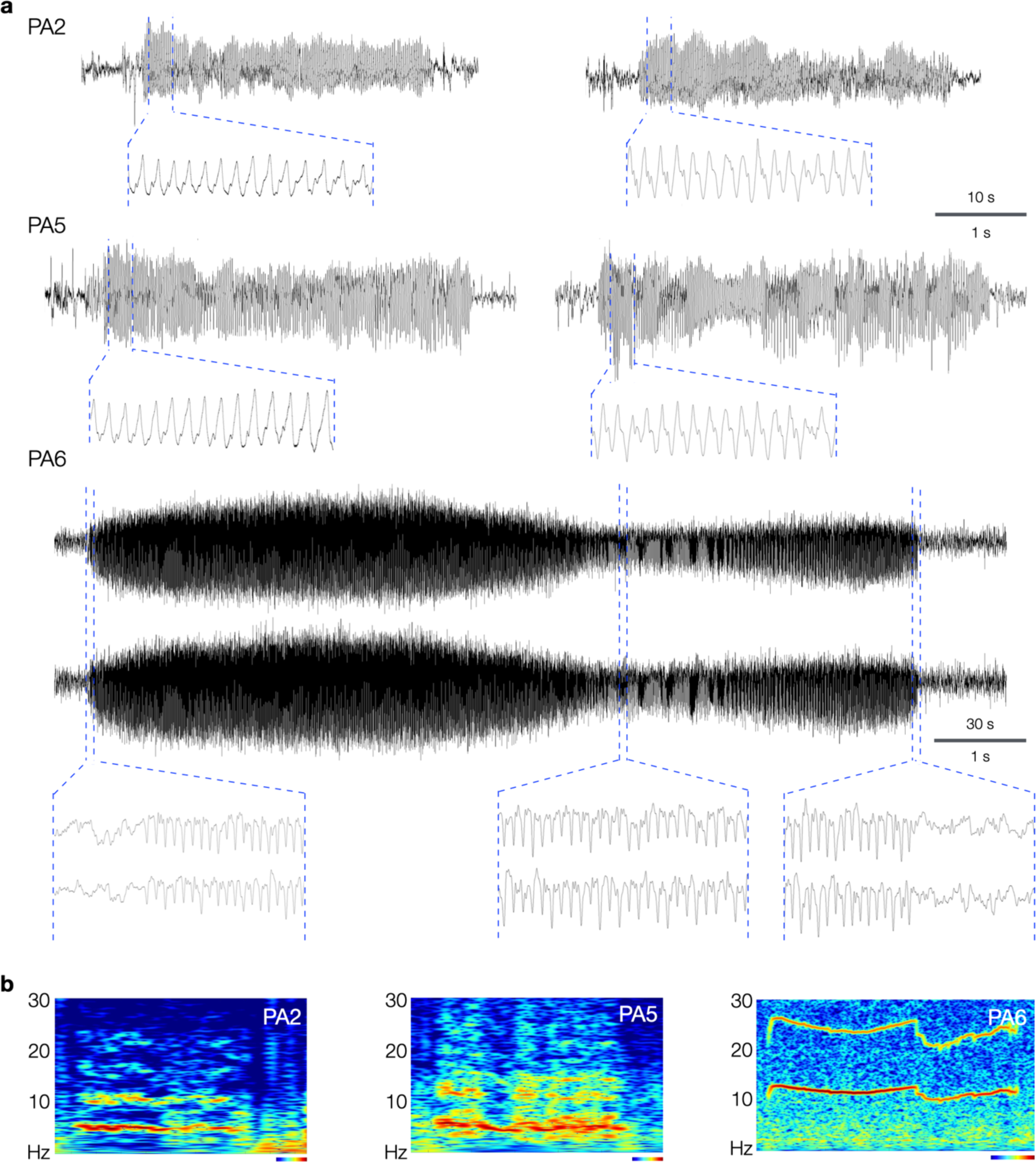
*Grin2a*-A243V PE3b-treated mice develop seizures with spike-and-wave morphology. **a**, Representative EEG traces from 3 PE-electroporated animals (PA2, PA5, and PA6) that developed spontaneous seizures after in utero prime editing with *Grin2a*-A243V. PA2 and PA5 developed frequent focal seizures, while PA6 presented a generalized seizure (EEG traces of both hemispheres shown) and sparse epileptiform events. Insets display magnifications of the indicated positions, showing spike-and-wave morphologies. Time-scale bars as indicated for full traces (top values) and insets (bottom values). **b**, Morlet wavelet spectrograms of seizures in (a) showing characteristic dominant frequency bands and harmonics. Heatmap and time-scale bars = 10 s (PA2 and PA5) and 30 s (PA6).

Two of the PA animals (PA2 and PA5) showed frequent, spontaneous motor seizures associated with asymmetric tonic posturing with hemiclonic movements (Supplementary Videos S1-4). Focal motor and secondarily generalized seizures are both typical of patients with SeLECTs. As shown in the representative traces in Fig. 4, events in PA2 and PA5 were electrographically characterized by sharply contoured, evolving spike-and-wave discharges in the ∼5 Hz range. As evident in example spectrograms (Fig 4b) and averaged traces (Fig. 5a and Extended Data Fig. 4), these events have discrete onset and termination with consistent frequencies. This event type represented the majority of observed seizures. A second seizure type was observed in animal PA6, a single generalized electrographic seizure without a motor component occurring during sleep (Fig. 4, PA6).

**Fig. 5.**
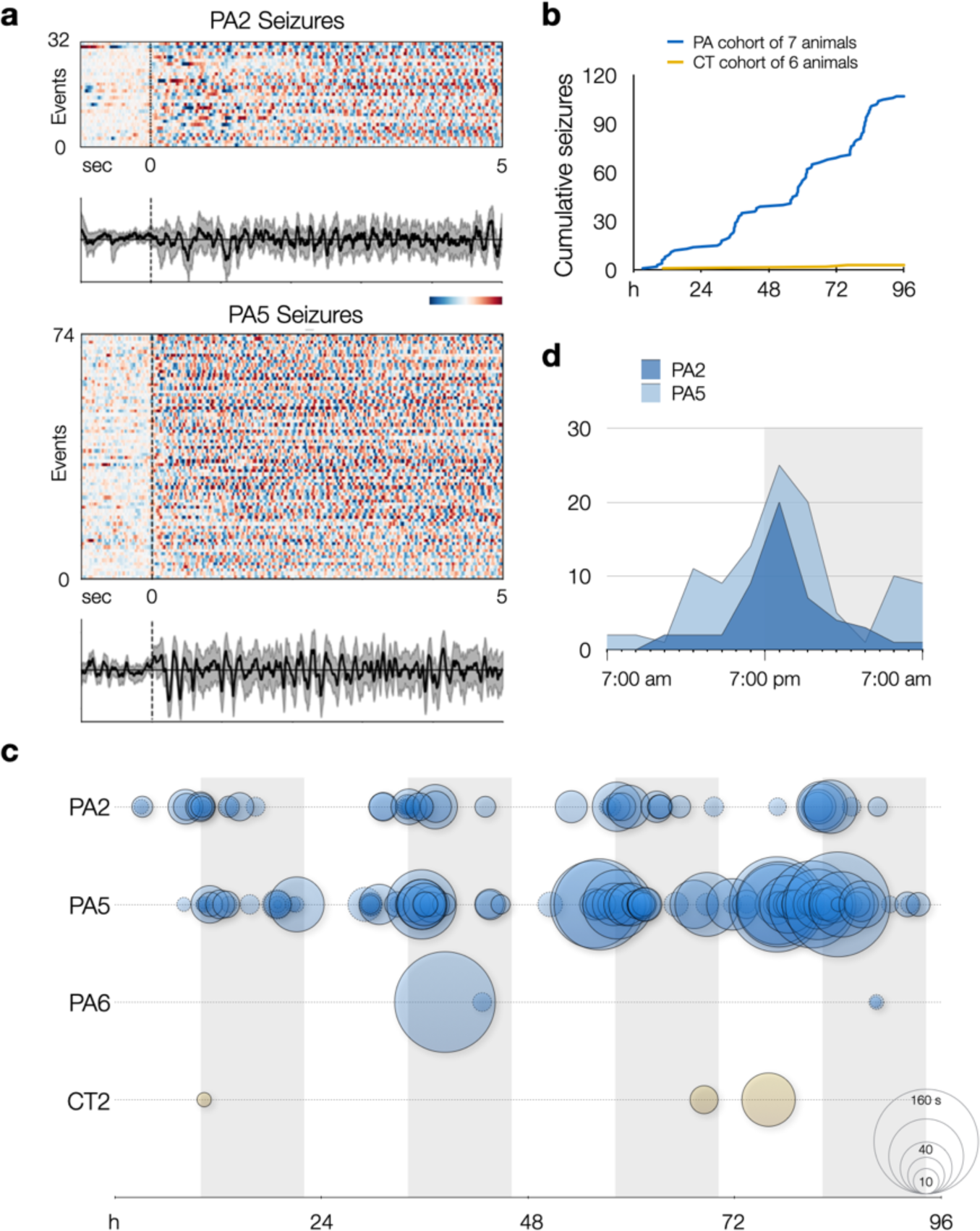
PE-electroporated PegAssist *Grin2a*-A243V mice have frequent spontaneous seizures with circadian patterns. a, Epoch charts and averaged traces show the stereotyped nature of seizures and epileptiform events. **a**, PA2 and PA5 epoch charts and averaged traces for events classified as seizures. Each row represents a separate event aligned at onset of event (t = 0) and plotted from t = −1 s to t = 5 s. Heatmap bar indicates 1 sec and standardized EEG amplitude from peak negative (dark blue) to peak positive (dark red). Traces show an averaged EEG signal in black with bootstrapped 95% confidence intervals in grey. **b**, Cumulative histogram of seizures over a 4-day recording period from PE-electroporated (PA; N=7) and control (CT; N=6) animals. Steps in cumulative histogram of PA cohort suggest circadian periodicity of seizures. See Extended Data Fig. 4 for analysis including epileptiform events. **c**, Seizures (solid circles) and epileptiform events (dashed circles) for each animal plotted by time and duration over 4-day recording period. Circle size indicates event duration as indicated on the bottom right reference circles (10, 20, 40, 80, 160 s). Grey vertical bars indicate daily dark cycle. **d**, Circadian histogram of seizures by hour in animal PA2 (dark blue) and PA5 (light blue). Seizures cluster in the 7:00-8:00 pm interval, corresponding to the beginning of the dark cycle when mice typically awaken.

Aggregating events within groups after blinded review of video and EEG recordings over 4 days, the PA cohort had a total of 107 seizures and an additional 56 epileptiform events, compared to 3 total events classified as seizures from one animal (CT3) electroporated with Cas9 and scrambled gRNA from the control cohort (Fig. 5 and Supplementary Table S1). We anticipated that only a subset of PA animals would manifest phenotypes due to the known variability of electroporation between individually treated embryos. For the PegAssist workflow, we propose that the treated animal cohort be segmented into spontaneously symptomatic and non-symptomatic animals. Symptomatic animals would then be monitored to establish individual baseline seizure frequency, as shown in Fig. 5, and would then each constitute a personalized patient model for use in N-of-1 type testing of compounds to assess antiepileptic efficacy.

Within each animal, seizures were highly stereotyped, both behaviorally (Supplementary Videos S1-4) and electrographically (Fig. 5). Averaged traces of all events demonstrate that each animal’s seizures displayed characteristic morphology and frequency (Extended Data Fig. 4), theoretically facilitating rapid automated analysis of seizure burden in subsequent N-of-1 trials. A further interesting pattern emerged when analyzing event distribution. In the two animals with frequent seizures, events displayed clear circadian rhythmicity (Fig. 5b), with seizures clustering around lights-off (Fig. 5c-d), the time when mice typically transition to periods of wakefulness^25,26^. This distribution mirrors a characteristic pattern in SeLECTS, wherein seizures most often occur during non-REM sleep or immediately after waking^27^, further adding clinical validity as a patient model. Finally, the presence of frequent spontaneous seizures contributes to the model’s utility in assessing patient-specific anti-seizure pharmacotherapy.

## Discussion

Genome editing has rapidly advanced biomedical research and is finding its first applications as therapeutics^28^. To overcome the low precision of most editors, current approaches to edit animals focus on ex vivo editing of stem cells or zygotes, which are then screened, clonally expanded, and bred into animal lines for further use. Here, we have taken a different approach to identify genome editing agents that are precise enough for direct in vivo use (Fig. 2) in order to circumvent the need for screening and breeding. We found prime editing with PE3b to genome edit neurons in vivo with high precision (Fig. 3).

Since its first description in 2019^14^, prime editing has been effectively used ex vivo on zygotes to generate edited animals through germline transmission^29–32^. While technically challenging, the potential for direct somatic cell prime editing in vivo would, in principle, allow for the generation of animal models without the labor-intensive and time-consuming process of germline transmission, which requires multiple rounds of breeding. Such prime editing directly in vivo was recently been demonstrated to be feasible in hepatocytes, retinal cells, lung epithelia, astrocytes, and cardiomyocytes^33–41^. Despite this potential, a disease model producing a clinical phenotype using direct in vivo prime editing in wild-type animals has not yet emerged.

Our results report the first demonstration of in vivo prime edited neurons and the first use of somatic cell prime editing to recapitulate a neurological patient phenotype. These data provide evidence for the feasibility, validity, and utility of in utero prime editing to rapidly model epilepsy patients in wild-type mice.

Exome sequencing is already proving clinically useful over broad epilepsy patient populations^42,43^. Combining genetic diagnosis established by patient exome sequencing with the ability we describe here to model individual variants in mice within clinically-useful timescales can provide a systematic path toward tailoring treatment options for genetic epilepsies.

We demonstrate proof-of-principle of this process by modeling an epilepsy patient carrying a variant of uncertain significance. The GRIN2A A243V point-variant was detected in exome sequencing, however no further information, such as pedigree or parent sequences, was available to assess whether the variant was causal to the pathology^18^. The uncertain pathogenic significance of the variant, a common attribution in clinical genetics^44^, makes it a poor candidate for traditional mouse modeling. The fact that prime editor-treated mice developed seizures with salient features of the pathology i) confirms the causal nature of the specific *GRIN2A* patient variant and ii) provides a ready animal model against which to test treatment options for variant-specific efficacy. The considerable genotypic and phenotypic diversity among patients with GRIN2-related disorders^45^ further highlights the importance of patient-specific animal models^4,46^.

The PegAssist approach holds several advantages over other modeling strategies: 1) The use of outbred wild-type animals diminishes cost and time of animal production, and increases genetic and behavioral robustness^47,48^. 2) Since editing in each cell is a distinct event, rare off-target edits are not amplified and are unlikely to influence outcomes, avoiding clonal artifacts that afflict animal-lines^49^. 3) The technology used is not species-limiting, meaning a similar approach can be used in non-rodent mammals, including non-human primates.

Prime editing has been successfully applied to a variety of genomic loci in vitro and more recently in somatic cell editing in vivo^33–41^, suggesting this approach is likely applicable to a wide range of genetic conditions, including diverse genetic epilepsies. We propose that this pipeline may be a valuable tool for assessing personalized pharmacotherapy options for individual patients, and for in vivo preclinical screens for drugs against ultra-rare and neglected genetic disease. The continued integration of cost-effective and rapid in vivo genome editing approaches to the field of precision medicine has great potential for further innovative clinical applications, extending the benefit of these new technologies to broad patient populations.

## Methods

### pegRNA design and PegAssist application

To facilitate the use and broad adoption of prime editing, we developed PegAssist and an accompanying webtool and set of plasmids for the design and production of custom Prime editing reagents. The python-based web application pegassist.app accepts input of target sequence and desired edits to produce pegRNA sequences and one-step cloning strategies based on the PegAssist plasmid set. The PegAssist platform offers PE2, PE3 and PE3b variant strategies, and further allows custom modifications of spacers and PAM sequences for versatility and use with further developments in genome editors (Extended Data Fig. 1).

The PegAssist python source code is available on github.com/pegassist. A graphical user interface was created using Heroku to compile a webtool available at pegassist.app.

To generate *Grin2a*-A243V pegRNAs, a custom spacer (20 nt) was first designed to minimize the distance between the protospacer adjacent motif (PAM) site and the edit position. The custom spacer and desired edit were used to generate automated reverse transcriptase template and primer binding sequence options and secondary nicking (PE3/PE3b) guides following the instructions outlined by the webtool. Based on the current knowledge of generalizable design rules, preference should be given to designs with a PBS length near 12nt with >30% GC content and a RT length near 14nt^17^.

### Plasmids

Double-stranded DNA oligonucleotides containing pegRNAs flanked by BbsI recognition sites were synthesized by Twist Bioscience. Sense and antisense oligonucleotides for knockout gRNA or PE3b gRNA with overhangs for golden gate assembly were synthesized by Integrated Design Technologies. Cloning was performed as previously described^12^. Briefly, pegRNA oligonucleotides were sub-cloned into pCR Blunt II-TOPO backbone (ThermoFisher). A golden gate assembly (GGA) with BbsI was used to clone the pegRNA into a custom backbone containing a hU6 promotor. The PE3/PE3b or knockout gRNA sense and antisense oligonucleotides were annealed and cloned by GGA into a pJ2^12^ containing a hU6 promoter. The vectors containing pegRNA and PE3/PE3b guides were used as template for PCR using KAPA HiFi HotStart DNA Polymerase with 2x Master Mix (Roche) to amplify parts containing the U6 promoter and either the pegRNA or PE3/PE3b guide using primers with BsaI recognition sites. These parts were assembled in a final vector by golden gate assembly using NEB Golden Gate Assembly Kit according to manufacturer’s recommendations. The open reading frame of PE2 was extracted from pCMV-PE2 (Addgene plasmid #132775) by PCR with primers (prRR842 and prRR849). PE2 was introduced into a pJ2 backbone by GGA to construct the final plasmid pJ2.CAG<EGFP-2A-PE2 [Lab plasmid ID: TU516]. Plasmid pCAG<myr-tdTomato expressing myristoylated tdTomato was subcloned from Addgene plasmid #26771 and used as a bright fluorescence electroporation reporter. Plasmid sequences were confirmed by Sanger sequencing by GeneWiz. Details and sequences of plasmids and oligonucleotides used are listed in Supplementary Table S2.

### BFP-to-GFP conversion assay

BFP-to-GFP conversion assays were performed as previously described^12^. Briefly, a modified HEK-293 cell line with genomically-encoded BFP was a gift from the Corn lab^50^. Cells were maintained in DMEM with GlutaMAX (ThermoFisher Scientific) supplemented with 10% fetal bovine serum. Cells were plated at a density of 20,000-22,500 cells/cm^2^ in 24-well plates prior to transfection using polyethylenimine, linear, MW 25000 (Polysciences) at 1 mg/mL in diH_2_O, then mixed in a 3:1 ratio with 750 ng total DNA diluted in Opti-MEM per well.

Conversion of BFP to GFP was analyzed by flow cytometry using an LSRII cell analyzer with HRS (BD Biosciences). A 407 nm laser with a 405/50 emission filter was used to detect BFP, while a 488 nm laser with a 505 LP mirror and a 530/30 emission filter was used to measure GFP.

### Mice

Experimental protocols involving animals were approved by the University of Maryland Baltimore Institutional Animal Care and Use Committee. Pregnant, outbred CD1 mice were obtained from Charles River Laboratories. *In utero* electroporation was performed on embryonic mice ambiguous to considerations of sex. Mice were weaned at P21 and EEG/EMG recordings were performed at 3-8 months.

### In utero electroporation

Cortical layer II/III pyramidal neurons were targeted by performing this procedure *in utero* on embryonic day 14.5 as previously described^51,52^. Briefly, plasmid DNA was combined to a maximum concentration of 4 µg/µL with equal molar ratios of relevant plasmids (pegRNA/PE3b duplex, prime editor, and fluorescent reporter). Dams were anesthetized with isoflurane with thermal support. The abdomen was prepared for surgery by removing hair and sanitizing the incision site using betadine and 70% ethanol. An incision of the skin and muscle layer along the midline exposed the uterine horns. A glass micropipette was pulled (Narishige PC-100) and beveled (Narishige PCR-45). The micropipette was attached to an aspirator and used to inject the prepared DNA mixture into the right ventricle of developing fetuses. Immediately upon injection, a series of 4 x 50 ms square pulses of 35 V (NEPA21 electro-kinetic platinum tweezertrodes on a BTX ECM-830 electroporator) was used to introduce the DNA into neural progenitor cells lining the ventricle. In a typical surgery 3-6 pups were electroporated. Following electroporation, the uterine horn was returned to the abdominal cavity, and the muscle and skin layers were closed using monofilament nylon sutures (AngioTech). After birth, electroporated pups were screened at post-natal day 0 for fluorescence using a fluorescence stereoscope (Leica MZ10f with X-Cite Fire LED light source). Positively screened pups were returned to the dam.

### Neuron Fluorescence Activated Cell Sorting and Next-Generation Sequencing

In utero electroporated mice were deeply anesthetized using isofluorane and euthanized. The brain was removed and immediately moved to pre-cooled dissociation medium (20 mM glucose, 0.8 mM kynurenic acid, 0.05 mM APV, 50 U/ml penicillin, 0.05 mg/mL streptomycin, 0.09 M Na_2_SO_4_, 0.03 M K_2_SO_4_, 0.014 M MgCl_2_) on ice. Using a fluorescence stereoscope (Leica MZ10f with X-Cite Fire LED light source), the electroporated region was dissected and transferred to a new tube containing ice-cold dissociation medium. Dissociation medium was aspirated until 1 mL remained and an activated papain solution (1:1 papain [Worthington-Biochem] with 13.6 mM Cysteine-HCL, 0.002% β-mercaptoethanol, and 2.4 mM EDTA pH 8.0 in MilliQ water) was added and incubated at 37° C for 30 minutes. Papain solution was removed, and tissue was washed three times and resuspended in 500 µL fresh dissociation medium. Samples were triturated 2-4 times using flame-polished borosilicate pipettes. Cell suspensions were sorted at low flow rates using a Wolf Benchtop Cell Sorter (Nanocollect Biomedical, Inc.) using red fluorescence from pCAG<myr-tdTomato for gating, after confirming overlap with green fluorescent signal from co-expressed pJ2.CAG<EGFP-2A-PE2. Approximately 3,000 cells were collected per sample. Sorted cells were pelleted and stored at −80°C.

Cells were lysed and total RNA was extracted using an AllPrep DNA/RNA Micro Kit (Qiagen). A reverse transcription reaction using a First Strand cDNA Synthesis Kit (Millipore Sigma) was used to create cDNA from the isolated RNA. The region surrounding the intended edit was amplified by PCR using primers containing universal adaptor sequences (prCR419 and prCR420). These amplicons were submitted to the Institute for Genome Sciences at the University of Maryland School of Medicine for sequencing, where samples were quantified, barcoded and sequenced on an Illumina NextSeq 550 (Illumina) according to manufacturer settings. An average of 8.9 million reads per sample were analyzed. GRCm39 was used as reference genome. Sanger sequencing of the target region from CD1 mice used in experiments were consistent with the reference genome (data not shown). Sequencing results were analyzed using CRISPResso2^53^ under prime editing mode. Default CRISPResso2 parameters were applied (quantification window 10, nicking guide sequence defined, scaffold match length 1). A contiguous quantification window was produced encompassing both pegRNA and PE3b gRNA target sequences.

### Electrophysiology

Whole-cell currents were recorded at room temperature from mouse cortical neurons which were isolated at P0 from E15 in utero electroporated animals and cultured 30 days in vitro (DIV) in order to allow for expression of GluN2A, which is known to be developmentally regulated^54^. Only cells expressing the red fluorescent marker were patched. Electrodes were pulled from borosilicate glass capillaries that were fire-polished to a resistance of 3-4 MΩ and filled with intracellular solution (mM): 135 CsCl, 35 CsOH, 4 MgATP, 0.3 Na_2_GTP, 10 HEPES and 1 EGTA, adjusted to pH 7.4 with CsOH. Cells were perfused with extracellular solutions containing (mM): 140 NaCl, 2.5 KCl, 1.8 CaCl_2_, 0.1 glycine, 10 HEPBS, 10 tricine, adjusted to pH 7.4 (NaOH). 40 µM cyanquixaline (CNQX) and 1µM ifenprodil were added to external solution to block GluN2B and AMPA receptors respectively. For solutions containing zinc, free Zn^2+^ concentrations in 10 mM tricine-buffered solutions were calculated using Maxchelator software (Chris Patton) using a binding constant of 10^-5^ M as previously reported^22^ and adjusted for our conditions. The final free zinc concentration was chelated to 67 nM by adding 200 mM ZnCl_2_ and 10 mM tricine into the working extracellular solution^55^. Currents were recorded with an Axopatch 200B amplifier (Molecular Devices) and digitized using an iTC-18 (InstruTECH). Currents were low-pass filtered at 2 kHz and sampled at 10 kHz using custom MATLAB (MathWorks) scripts. Drugs and agonists were applied during the patch recordings by means of an eight-barrel pen-perfusion system, with minimal dead space. In all whole-cell experiments, the cells were clamped at −80 mV. Solution containing NMDA (100 µM), or NMDA with 67 nM free Zn^2+^ was applied to elicit the current, usually for 5 sec every 2 min, using motor-driven valves.

For HEK 293 cell recordings, cells were cultured on glass coverslips coated with poly-D-lysine and transfected via the calcium phosphate method^56^ with 4–8 μg of rat GluN1-1a and GluN2A or GluN2A (A243V), and co-transfected with 2 μg green fluorescent protein. Culture media was exchanged 3–5 h post-transfection, and cells were maintained 24–48 hours in DMEM supplemented with 2 mM Mg^2+^ to prevent NMDA receptor-mediated cell death. The GluN1-1a and GluN2A plasmids were a kind gift from Gabriela Popescu (University at Buffalo). The mutation A243V was introduced into the GluN2A plasmid using the QuickChange II XL kit from Agilent. All portions of the resulting construct that were subject to PCR were confirmed by DNA sequencing. Recording conditions were identical to those listed above for neurons.

### Continuous EEG/EMG recording

Synchronous EEG/EMG and video recording was performed using a tethered, 3-channel recording system from Pinnacle Technology Inc. Prefabricated EEG headmounts were implanted with screw electrodes 3 mm behind Bregma and an EMG lead was implanted in the trapezius muscle of mice under isofluorane anesthesia. Mounted electrodes were fixed using dental epoxy. After minimum 72 hours recovery, mice were placed in recording chambers for synchronous EEG/EMG and video recording for approximately 4 days.

The EEG/EMG data were exported and blinded before primary review. An initial investigator manually annotated possible epileptiform events and exported epochs for secondary review by a blinded expert (author PFD) and classification based on the American Clinical Neurophysiology Society’s most recent standardized criteria^57^. Specifically, seizure was defined as any rhythmic epileptiform discharge with either a) time-locked, consistent behavioral correlate, b) discharges averaging >2.5Hz for at least 10s, or c) with definite spatiotemporal evolution and lasting at least 10s. Based on accepted ACNS criteria for human EEG, the latter two categories would be termed “electrographic seizure” as opposed to “electroclinical seizure”, but in this analysis we did not make such a distinction.

EEG, EMG, and video data were used to visually identify artifacts associated with movement. If movement associated with EEG change was stereotyped across events (within animals) (see videos S1-4), the abnormal movements associated with these events were considered clinical correlates. Seizure events were further classified as lateralized for events occurring in only one EEG channel or generalized for events with synchronous activity between both EEG channels. If similar activity was seen that was without clear clinical correlate, did not last 10s duration, and did not have clear spatiotemporal evolution, these events were classified as interictal discharges (“epileptiform” in Supplementary Table S1 and Extended Data Fig. 4) as opposed to seizures. Seizure epoch charts were generated from annotated seizure bouts using MNE-Python following standardization using the SciKit StandardScaler function. Seizure frequency was calculated as the total number of events divided by days of recording.

### Tissue Fixation and Immunolabeling

Tissue was fixed by transcardial perfusion of mice using PBS and 4% paraformaldehyde with 24 hours post-fixation in 4% paraformaldehyde at 4°C. Tissue slices were cut to 80 µm using a vibrating microtome (Leica). For immunolabeling, slices were incubated in a blocking solution of 5% BSA, 0.3% TritonX-100, and 0.05% sodium azide in PBS for 2 hours while rocking at room temperature. Primary antibodies were diluted 1:1000 in blocking solution, and tissue was incubated overnight, rocking at room temperature. Slices were washed 3x 30 minutes in PBS while rocking, then incubated with secondary antibody at 1:1000 in PBS at room temperature for 4 hours. After 3x 30-minute PBS washes the slices were mounted on slides with either Fluoromount-G Mounting Medium with or without DAPI (ThermoFisher Scientific).

### Microscopy

Fluorescence images were acquired using a Nikon Ti2-E inverted epifluorescence microscope. Images were analyzed using NIS Elements (Nikon). Proximal z-stacks were acquired using a 10x objective, then extended depth of focus and stitching were used to compile a single slice image.

## Acknowledgements

We are grateful to Gabriela Popescu (Buffalo), Linda Nowak (Ithaca), and Cornelia Poulopoulou (Athens) for helpful discussions and technical advice. We thank Elise Caraker, Corinne Martin, Robert Lease (Baltimore), and Corey Flynn (Boston) for technical expertise. We are grateful for the technical and instrumentation support of the University of Maryland School of Medicine Center for Innovative Biomedical Resources through the Confocal Imaging Facility, the Flow Cytometry Facility, and the Institute for Genome Sciences Genomics Facility. This work was made possible through funding support from the US National Institutes of Health through DP2MH122398 (AP), R01HL149926 (IED), R37NS125632 (PBC). CDR was funded by the Training Program in Integrative Membrane Biology through T32GM008181; RRR was funded by the Training Grant in Cancer Biology through T32CA154274; DCOV was funded by a Postdoctoral Fellowship by the American Heart Association through; SBEW was funded by a Klingenstein-Simons Fellowship Award in Neurosciences by the Esther A. & Joseph Klingenstein Fund and the Simons Foundation.

---

**Extended Data Fig. 1.**
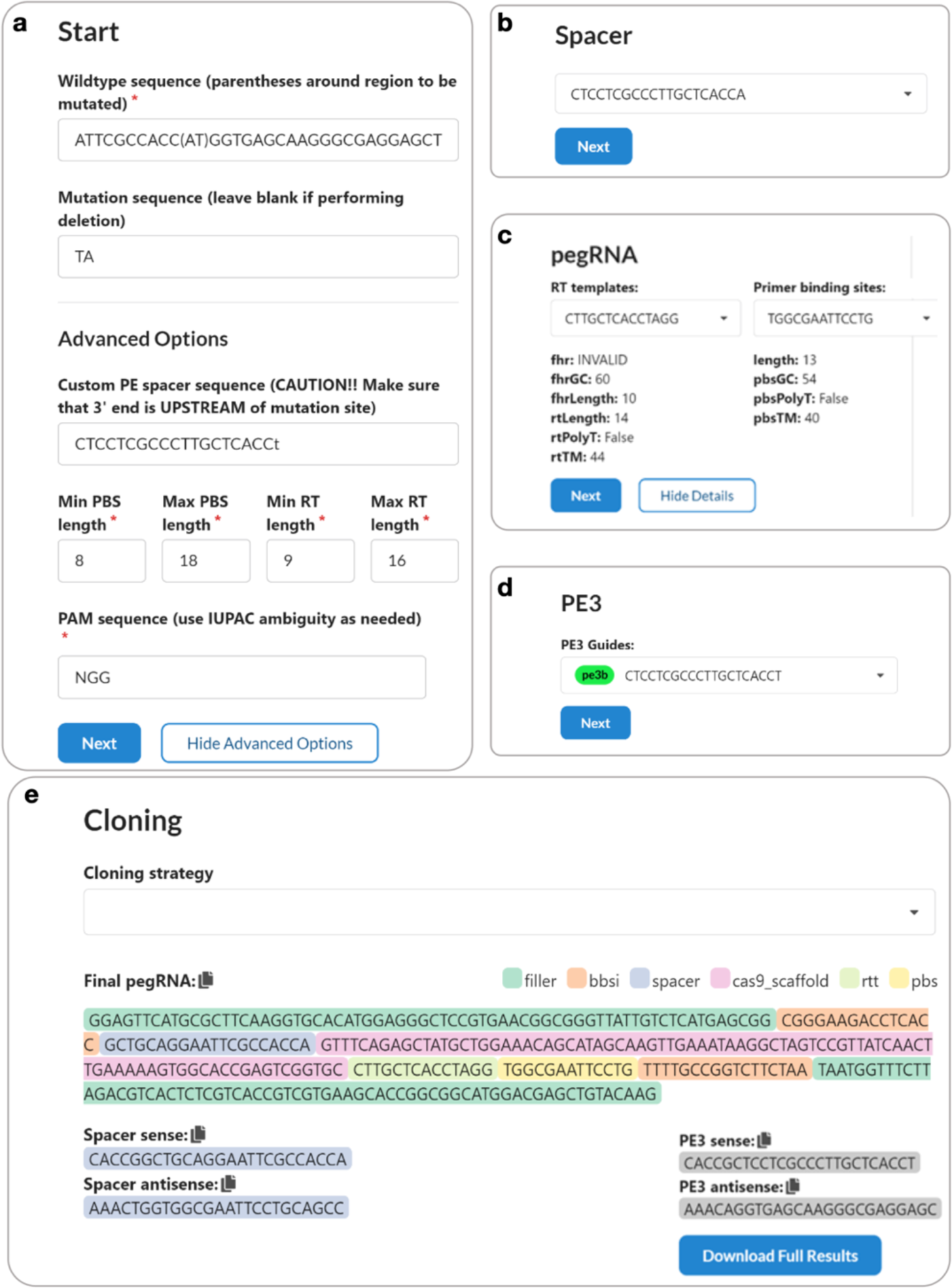
Pegassist.app for pegRNA and PE3b guide design. **a**, pegassist.app accepts input of a WT sequence with the desired edit in parentheses. The edited sequence is entered, and users can select from additional options including input of a pre-designed spacer sequence. **b**, A spacer sequence is selected from the dropdown menu. **c**, RT template and Primer binding site options are presented with warnings against low efficiency. **d**, All PE3/PE3b options are presented for selection. **e**, pegassist.app displays a final pegRNA and the required oligos based on the selected cloning method.

**Extended Data Fig. 2.**
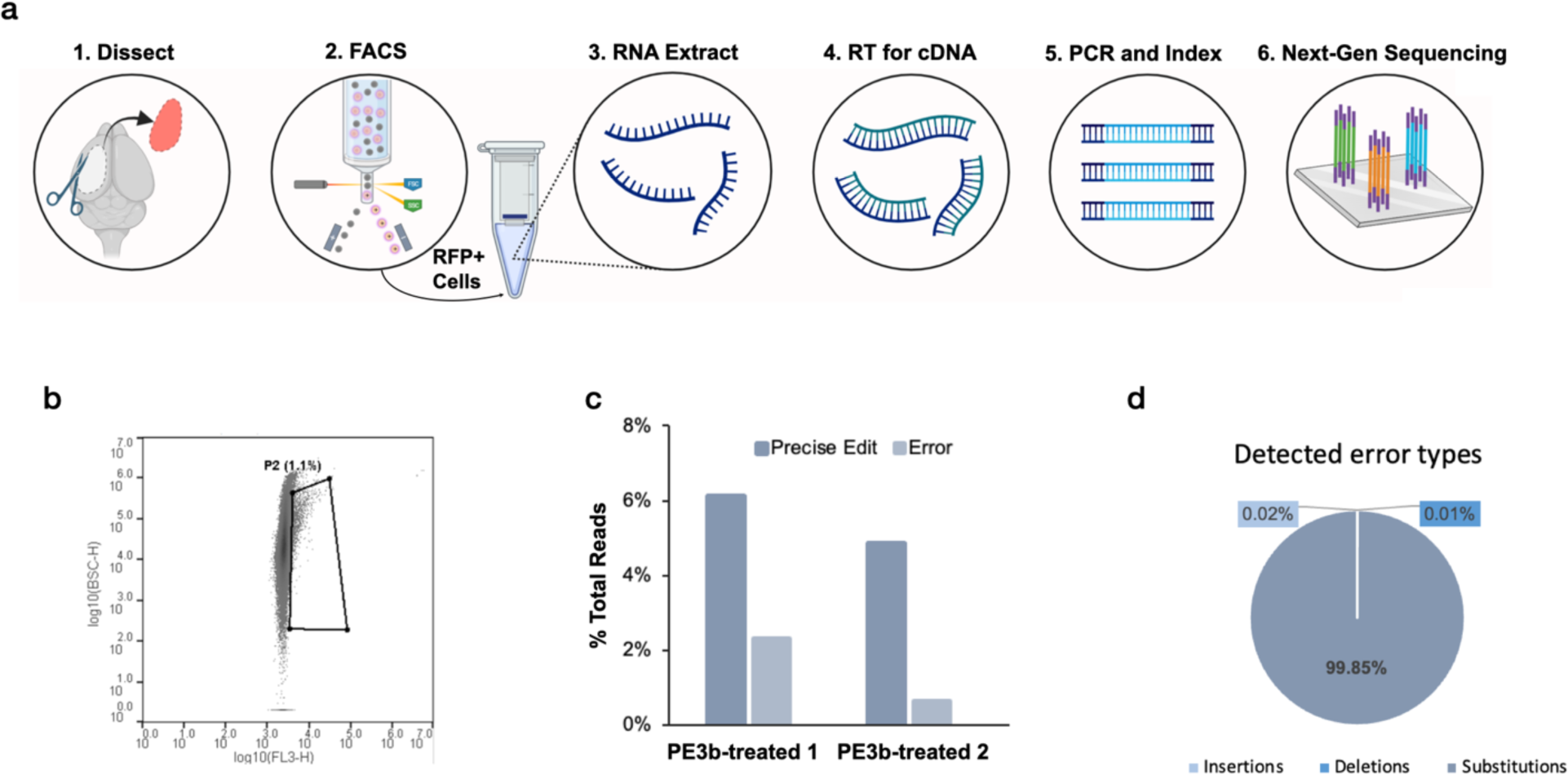
In vivo prime editing of *Grin2a* is precise and causes few indels. **a**, Schematic representation of the sequencing workflow. Following in utero electroporation with PE3b agents, the PE-expressing region of cortex was dissected from adult mice. Cells were triturated and sorted for red fluorescence. RNA was extracted to generate cDNA by reverse transcription. Target sequence was amplified by PCR. Amplicons were indexed and sequenced on the Illumina NextSeq platform. **b**, FACS plot of neurons gated to collect electroporated red fluorescent neurons (∼1% of total neurons). y axis plots side scatter and x axis plots red fluorescence intensity. **c**, Percentage of reads with either the intended edit or any error in the two PE3b-treated animals sequenced. The intended edit is detected approximately 5 times more frequently than the aggregate of all proximal erroneous edits. d, Breakdown of error types detected in PE3b-treated samples. Most errors are substitutions and may be expected to have less deleterious effects on the resulting protein than indels.

**Extended Data Fig. 3.**
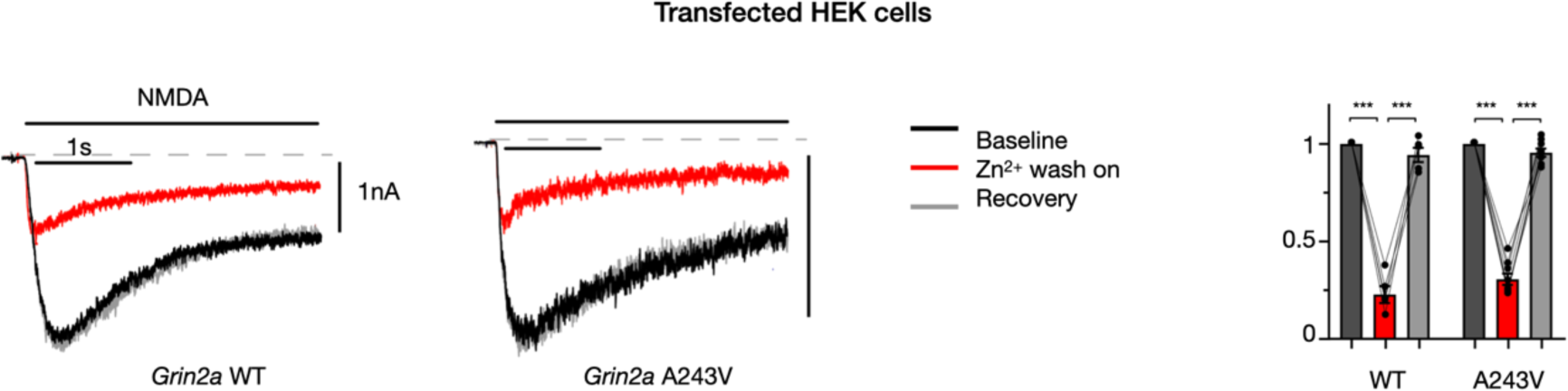
A243V in *Grin2a* does not affect Zn^2+^ blockade of NMDA currents. Example traces and amplitudes of currents recorded from HEK cells co-transfected with *GRIN1* and either WT (n=5) or A243V (n=8) variant of *GRIN2A*; baseline (black), Zn^2+^ wash-on (red), recovery (grey); amplitudes normalized to baseline of each sample. A243V does not change the sensitivity to Zn^2+^ blockade of NMDA currents (*p<0.05, ***p<0.001, via students T-test).

**Extended Data Fig. 4.**
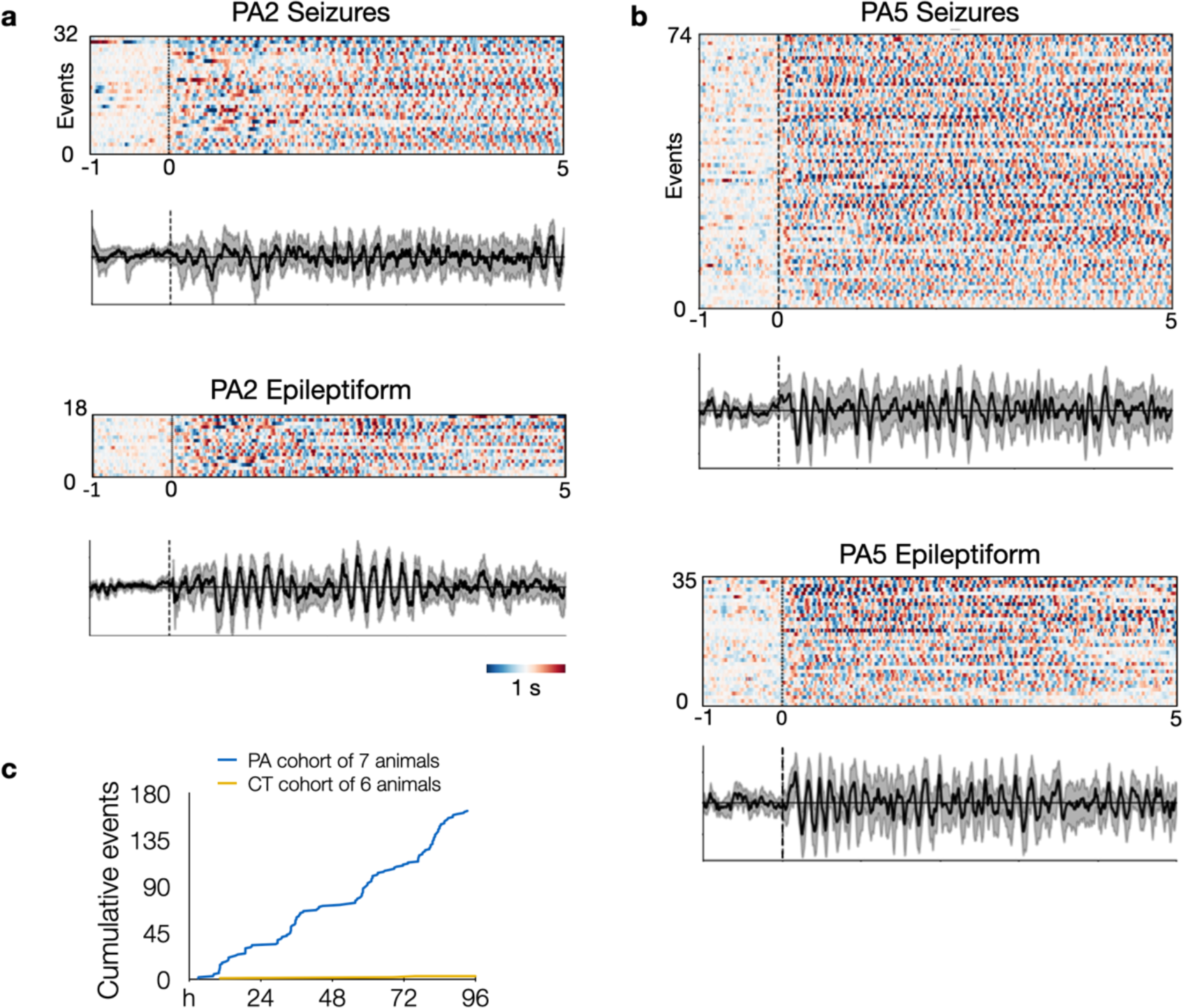
Epoch charts and averaged traces show the stereotyped nature of seizures and epileptiform events. **a**, PA2 and PA5 epoch charts and averaged traces for events classified as seizures (same as in Fig. 5a) or epileptiform events. Each row represents a separate event aligned at onset of event (t = 0) and plotted from t = −1 s to t = 5 s. Heatmap bar indicates 1 sec and standardized EEG amplitude from peak negative (dark blue) to peak positive (dark red). Traces show an averaged EEG signal in black with bootstrapped 95% confidence intervals in grey. **c**, Pooled cumulative events (including both seizures and epileptiform events) between PA and CT groups over 4 days of EEG recording.

**Supplementary Table S1.** Individual annotation of seizures and epileptiform events (“Events” tab), and details of PA and CT animal cohorts (“Cohorts” tab).

**Supplementary Table S2.** Plasmids and oligonucleotides used: identifiers, purpose, and sequences.

**Supplementary Videos S1-2.** Videos of seizures corresponding to the two example traces from animal PA2 in Fig. 4a.

**Supplementary Videos S3-4.** Videos of seizures corresponding to the two example traces from animal PA5 in Fig. 4a.

